# Meta-analysis of exome array data identifies six novel genetic loci for lung function

**DOI:** 10.1101/164426

**Authors:** Victoria E Jackson, Jeanne C Latourelle, Louise V Wain, Albert V. Smith, Megan L. Grove, Traci M Bartz, Maen Obeidat, Michael A Province, Wei Gao, Beenish Qaiser, David J Porteous, Patricia A Cassano, Tarunveer S. Ahluwalia, Niels Grarup, Jin Li, Elisabeth Altmaier, Jonathan Marten, Sarah E Harris, Ani Manichaikul, Tess D Pottinger, Ruifang Li-Gao, Allan Lind-Thomsen, Anubha Mahajan, Lies Lahousse, Medea Imboden, Alexander Teumer, Bram Prins, Leo-Pekka Lyytikäinen, Gudny Eiriksdottir, Nora Franceschini, Colleen M Sitlani, Jennifer A Brody, Yohan Bossé, Wim Timens, Aldi Kraja, Anu Loukola, Wenbo Tang, Yongmei Liu, Jette Bork-Jensen, Johanne M. Justesen, Allan Linneberg, Leslie A. Lange, Rajesh Rawal, Stefan Karrasch, Jennifer E Huffman, Blair H Smith, Gail Davies, Kristin M Burkart, Josyf C Mychaleckyj, Tobias N. Bonten, Stefan Enroth, Lars Lind, Guy G Brusselle, Ashish Kumar, Beate Stubbe, Mika Kähönen, Annah B. Wyss, Bruce M Psaty, Susan R Heckbert, Ke Hao, Taina Rantanen, Stephen B Kritchevsky, Kurt Lohman, Tea Skaaby, Charlotta Pisinger, Torben Hansen, Holger Schulz, Ozren Polasek, Archie Campbell, John M Starr, Stephen S Rich, Dennis O. Mook-Kanamori, Åsa Johansson, Erik Ingelsson, André G Uitterlinden, Stefan Weiss, Olli T. Raitakari, Vilmundur Gudnason, Kari E. North, Sina A Gharib, Don D Sin, Kent D Taylor, George T O’Connor, Jaakko Kaprio, Tamara B Harris, Oluf Pedersen, Henrik Vestergaard, James G. Wilson, Konstantin Strauch, Caroline Hayward, Shona Kerr, Ian J Deary, R Graham Barr, Renée de Mutsert, Ulf Gyllensten, Andrew P Morris, M. Arfan Ikram, Nicole M Probst-Hensch, Sven Gläser, Eleftheria Zeggini, Terho Lehtimäki, David P Strachan, Josee Dupuis, Alanna C. Morrison, Ian P Hall, Martin D Tobin, Stephanie J London

**Affiliations:** Department of Health Sciences, University of Leicester, Leicester, UK; Department of Neurology, Boston University school of Medicine, Boston MA; National Institute for Health Research, Leicester Respiratory Biomedical Research Unit, Glenfield Hospital, Leicester, UK; Icelandic Heart Association, 201 Kopavogur, Iceland; University of Iceland, 101 Reykjavik Iceland; Human Genetics Center, Department of Epidemiology, Human Genetics, and Environmental Sciences, School of Public Health, The University of Texas Health Science Center at Houston. Houston TX 77030; Cardiovascular Health Research Unit, Departments of Medicine and Biostatistics, University of Washington, Seattle, WA, USA, 98101; The University of British Columbia Centre for Heart Lung Innovation, St Paul’s Hospital, Vancouver, BC, Canada; Department of Genetics, Washington University School of Medicine, St. Louis, MO.; Department of Biostatistics, Boston University School of Public Health, Boston, MA, USA; Institute for Molecular Medicine Finland (FIMM), University of Helsinki, FI-00014, Helsinki, Finland; Centre for Genomic & Experimental Medicine, MRC Institute of Genetics & Molecular Medicine, University of Edinburgh, Edinburgh, EH4 2XU, UK; Division of Nutritional Sciences, Cornell University, Ithaca NY; Department of Healthcare Policy and Research, Division of Biostatistics and Epidemiology, Weill Cornell Medical College NY, NY; The Novo Nordisk Foundation Center for Basic Metabolic Research, Faculty of Health and Medical Sciences, University of Copenhagen, 2200 Copenhagen, Denmark; Steno Diabetes Center Copenhagen, Gentofte, Denmark; Department of Medicine, Division of Cardiovascular Medicine, Stanford University School of Medicine, Palo Alto, CA, USA; Research Unit of Molecular Epidemiology, Institute of Epidemiology II, Helmholtz Zentrum München, German Research Center for Environmental Health, 85764 Neuherberg, Germany; Medical Research Council Human Genetics Unit, Institute of Genetics and Molecular Medicine, University of Edinburgh, Edinburgh EH4 2XU, UK; Centre for Cognitive Ageing and Cognitive Epidemiology, University of Edinburgh, Edinburgh, UK, EH8 9JZ; Centre for Genomic and Experimental Medicine, University of Edinburgh, Edinburgh, UK, EH4 2XU; Center for Public Health Genomics, Department of Public Health Sciences, University of Virginia, Charlottesville, VA; Department of Medicine, College of Physicians and Surgeons, Columbia University, New York, NY; Department of Preventive Medicine - Division of Health and Biomedical Informatics, Northwestern University - Feinberg School of Medicine, Chicago, IL; Department of Clinical Epidemiology, Leiden University Medical Center, Leiden, the Netherlands, 2333 ZA; Department of Immunology, Genetics, and Pathology, Biomedical Center, SciLifeLab Uppsala, Uppsala University, SE-75108 Uppsala, Sweden; Center for health and infectious disease research, Rigshospitalet, Copenhagen, Denmark; Wellcome Trust Centre for Human Genetics, University of Oxford, Oxford, UK; Respiratory Medicine, Ghent University Hospital, Ghent, Belgium, BE9000; Epidemiology, Erasmus Medical Center, Rotterdam, the Netherlands, 3000CA; Swiss Tropical and Public Health Institute, Basel, Switzerland; University of Basel, Switzerland; Institute for Community Medicine, University Medicine Greifswald, 17475 Greifswald, Germany; Human Genetics, Wellcome Trust Sanger Institute, Hinxton, UK, CB10 1SA; Department of Clinical Chemistry, Fimlab Laboratories, Tampere 33520, Finland; Department of Clinical Chemistry, Faculty of Medicine and Life Sciences, University of Tampere, Tampere 33014, Finland; Department of Epidemiology, Gillings School of Global Public Health, University of North Carolina, Chapel Hill, NC 27514; Cardiovascular Health Research Unit, Department of Medicine, University of Washington, Seattle, WA, USA, 98101; Institut universitaire de cardiologie et de pneumologie de Québec, Department of Molecular Medicine, Laval University, Québec, Canada; University of Groningen, University Medical Center Groningen, Department of Pathology and Medical Biology, HPC-EA10, Groningen, The Netherlands, NL9713 GZ; University of Groningen, University Medical Center Groningen, Groningen Research Institute for Asthma and COPD; Boeringher Ingleheim, Danbury, CT; Wake Forest School of Medicine, Winston-Salem, North Carolina, USA; Research Centre for Prevention and Health, Capital Region of Denmark, 2600 Copenhagen, Denmark; Department of Clinical Experimental Research, Rigshospitalet, 2600 Glostrup, Denmark; Department of Clinical Medicine, Faculty of Health and Medical Sciences, University of Copenhagen, 2200 Copenhagen, Denmark; Department of Medicine, Division of Bioinformatics and Personalized Medicine, University of Colorado Denver, Aurora, CO, USA; Institute of Epidemiology I, Helmholtz Zentrum München, German Research Center for Environmental Health, 85764 Neuherberg, Germany; Institute and Outpatient Clinic for Occupational, Social and Environmental Medicine, Ludwig-Maximilians-Universität, Munich, Germany.; Division of Population Health Sciences, Ninewells Hospital and Medical School, University of Dundee, Dundee, DD1 9SY, UK; Psychology, University of Edinburgh, Edinburgh, UK, EH8 9JZ; Department of Pulmonology, Leiden University Medical Center, Leiden, the Netherlands, 2333 ZA; Department of Public Health and Primary Care, Leiden University Medical Center, Leiden, the Netherlands, 2333 ZA; Department of Medical Sciences, Uppsala University Hospital, Uppsala, Sweden; Respiratory Medicine, Erasmus Medical Center, Rotterdam, the Netherlands, 3000CA; Institute of Environmental Medicine, Karolinska Institutet, Stockholm, Sweden; Internal Medicine B, University Medicine Greifswald, 17475 Greifswald, Germany; Department of Clinical Physiology, Tampere University Hospital, Tampere 33521, Finland; Department of Clinical Physiology, Faculty of Medicine and Life Sciences, University of Tampere, Tampere 33014, Finland; Epidemiology Branch, National Institute of Environmental Health Sciences, National Institute of Health, Dept of Health and Human Services, Research Triangle Park, NC, 27709, USA; Cardiovascular Health Research Unit, Departments of Epidemiology, Medicine and Health Services, University of Washington, Seattle, WA, 98101; Kaiser Permanente Washington Health Research Insititute; Cardiovascular Health Research Unit, Department of Epidemiology, University of Washington, Seattle, WA, 98101; Department of Genetics and Genomic Sciences, Icahn School of Medicine at Mount Sinai, One Gustave L. Levy Place, New York, NY 10029-6574; Icahn Institute of Genomics and Multiscale Biology, Icahn School of Medicine at Mount Sinai, One Gustave L. Levy Place, New York, NY 10029-6574; Department of Health Sciences, University of Jyväskylä FI-40014 Jyväskylä, Finland; Sticht Center on Aging, Wake Forest School of Medicine, Winston-Salem, North Carolina; Comprehensive Pneumology Center Munich (CPC-M), Member of the German Center for Lung Research, Munich, Germany; Faculty of Medicine, University of Split, Split, Croatia; Alzheimer Scotland Research Centre, University of Edinburgh, Edinburgh, UK, EH8 9JZ; Department of Medical Sciences, Molecular Epidemiology and Science for Life Laboratory, Uppsala University, Uppsala, Sweden; Department of Medicine, Division of Cardiovascular Medicine, Stanford University School of Medicine, CA 94305, USA; Internal Medicine, Erasmus Medical Center, Rotterdam, the Netherlands, 3000CA; Interfaculty Institute for Genetics and Functional Genomics, University Medicine Greifswald, 17475 Greifswald, Germany; DZHK (German Centre for Cardiovascular Research), partner site: Greifswald, Greifswald, Germany.; Department of Clinical Physiology and Nuclear Medicine, Turku University Hospital, Turku 20521, Finland; Research Centre of Applied and Preventive Cardiovascular Medicine, University of Turku, Turku 20014, Finland; Department of Epidemiology and Carolina Center for Genome Sciences, University of North Carolina, Chapel Hill, NC 27514; Computational Medicine Core, Center for Lung Biology, UW Medicine Sleep Center, Department of Medicine, University of Washington, Seattle, WA, 98109; Respiratory Division, Department of Medicine, University of British Columbia, Vancouver, BC, Canada; Institute for Translational Genomics and Population Sciences and Department of Pediatrics, Los Angeles Biomedical Research Institute at Harbor-UCLA Medical Center, Torrance, CA; Pulmonary Center, Department of Medicine, Boston University School of Medicine, Boston, MA 02118, USA; National Heart, Lung, and Blood Institute’s and Boston University’s Framingham Heart Study, Framingham, MA 01702, USA; Department of Public Health, University of Helsinki, FI-00014, Helsinki, Finland; Department of Health, National Institute for Health and Welfare, FI-00271 Helsinki, Finland; National Institute on Aging, National Institutes of Health, Bethesda, Maryland; Steno Diabetes Center Copenhagen, Gentofte, Denmark.; Department of Physiology and Biophysics, University of Mississippi Medical Center, Jackson, MS, USA; Institute of Genetic Epidemiology, Helmholtz Zentrum München, German Research Center for Environmental Health, 85764 Neuherberg, Germany; Institute of Medical Informatics, Biometry and Epidemiology, Chair of Genetic Epidemiology, Ludwig-Maximilians-Universität, 81377 Munich, Germany; Department of Epidemiology, Mailman School of Public Health, Columbia University, New York, NY; Department of Biostatistics, University of Liverpool, Liverpool, UK; Radiology, Erasmus Medical Center, Rotterdam, the Netherlands, 3000CA; Neurology, Erasmus Medical Center, Rotterdam, the Netherlands, 3000CA; Vivantes Klinikum Spandau Berlin, Department of Internal Medicine - Pulmonary Diseases, 13585 Berlin; Population Health Research Institute, St George’s, University of London, Cranmer Terrace, London SW17 0RE, UK; Division of Respiratory Medicine, University of Nottingham, Nottingham, UK

## Abstract

Over 90 regions of the genome have been associated with lung function to date, many of which have also been implicated in chronic obstructive pulmonary disease (COPD). We carried out meta-analyses of exome array data and three lung function measures: forced expiratory volume in one second (FEV1), forced vital capacity (FVC) and the ratio of FEV1 to FVC (FEV1/FVC). These analyses by the SpiroMeta and CHARGE consortia included 60,749 individuals of European ancestry from 23 studies, and 7,721 individuals of African Ancestry from 5 studies in the discovery stage, with follow-up in up to 111,556 independent individuals. We identified significant (P<2·8x10^-7^) associations with six SNPs: a nonsynonymous variant in *RPAP1*, which is predicted to be damaging, three intronic SNPs (*SEC24C, CASC17* and *UQCC1*) and two intergenic SNPs near to *LY86* and *FGF10*. eQTL analyses found evidence for regulation of gene expression at three signals and implicated several genes including *TYRO3* and *PLAU*. Further interrogation of these loci could provide greater understanding of the determinants of lung function and pulmonary disease.

## Introduction

Lung function measures are important predictors of mortality and morbidity and form the basis for the diagnosis of chronic obstructive pulmonary disease (COPD), a leading cause of death globally.^1^ Lung function is largely affected by environmental factors such as smoking and exposure to air pollution; however there is also a genetic component, with heritability estimates ranging between 39-66%.^2–5^ A number of large-scale genome-wide association studies (GWAS) of lung function have successfully identified single nucleotide polymorphisms (SNPs) influencing lung function at over 90 independent loci.^6–13^ Associations have also been identified in GWAS of COPD;^14–18^ however the identification of disease associated SNPs has been restricted by limited sample sizes. Many signals first identified in powerful studies of quantitative lung function traits, have been found to be associated with risk of COPD, highlighting the potential clinical usefulness of comprehensive identification of lung function associated SNPs.^13^

Low frequency (minor allele frequency (MAF) 1-5%) and rare (MAF<1%) variants, have been largely underexplored by GWAS to date. Exome arrays have been designed to facilitate the investigation of these low frequency and rare variants, predominately within coding regions, in large sample sizes. Alongside a core content of rare coding SNPs, the exome array additionally includes common variation including tags for previously identified GWAS hits, ancestry informative SNPs, a grid of markers for estimating identity by descent and a random selection of synonymous SNPs.^19^

## Results

We carried out a meta-analysis of exome array data and three lung function measures: forced expiratory volume in one second (FEV1), forced vital capacity (FVC) and the ratio of FEV1 to FVC (FEV1/FVC). These analyses included 68,470 individuals from the SpiroMeta and CHARGE consortia in a discovery analysis, with follow-up in an independent sample of up to 111,556 individuals. All studies are listed with their study-specific sample characteristics in Table 1, with full study descriptions, including details of spirometry and other measurements described in the Supplementary Note. The genotype calling procedures implemented by each study (Supplementary Table 1) and quality control of genotype data are described in the Supplementary Methods. We have undertaken both single variant analyses, and gene-based associations, which test for the joint effect of several rare variants in a gene (see methods for details).

**Table 1.** Sample characteristics of 11 SpiroMeta and 12 CHARGE studies contributing to the discovery analyses and 3 studies contributing to the replication analyses.

| Discovery Studies |  |  |  |  |  |  |  |
| --- | --- | --- | --- | --- | --- | --- | --- |
| SpiroMeta Studies | Total Sample | n (%) Male | Ever Smokers, n (%) | Age, mean (SD) | FEV<sub>1</sub>, litres. mean (SD) | FVC, litres. mean (SD) | FEV<sub>1</sub>/FVC, mean (SD) |
| 1958 British Birth Cohort (B58C) | 5270 | 2961 (56.2%) | 2866 (53.3%) | 44.00 (0.00) | 3.35 (0.79) | 4.29 (1.03) | 0.788 (0.09) |
| Generation Scotland (GS:SFHS) | 8164 | 3413 (41.8%) | 3806 (46.6%) | 51.59 (13.33) | 2.78 (0.87) | 3.91 (1.01) | 0.710 (0.12) |
| Cooperative Health Research in the Region of Augsburg (KORA F4) | 1447 | 701 (48.5%) | 900 (62.2%) | 54.82 (9.66) | 3.24 (0.85) | 4.20 (1.04) | 0.771 (0.07) |
| CROATIA-Korcula cohort (KORCULA) | 791 | 296 (36.8%) | 418 (52.0%) | 55.56 (13.69) | 2.72 (0.83) | 3.29 (0.95) | 0.829 (0.10) |
| Lothian Birth Cohort 1936 (LBC1936) | 974 | 501 (50.6%) | 554 (55.9%) | 69.55 (0.84) | 2.38 (0.67) | 3.04 (0.87) | 0.787 (0.10) |
| Study of Health in Pomerania (SHIP) | 1681 | 831 (49.4%) | 955 (56.8%) | 52.25 (13.43) | 3.29 (0.88) | 3.88 (1.03) | 0.848 (0.07) |
| Northern Swedish Population Health Study (NSPHS) | 880 | 407 (46.3%) | 122 (13.9%) | 49.13 (19.96) | 2.93 (0.90) | 3.53 (1.06) | 0.831 (0.09) |
| Prospective Investigation of the Vasculature in Uppsala Seniors (Pivus) | 836 | 413 (49.4%) | 426 (51.0%) | 70.20 (0.17) | 2.44 (0.68) | 3.20 (0.87) | 0.76 (0.10) |
| Swiss study on Air Pollution and Lung Disease in adults (SAPALDIA) | 2707 | 1379 (50.9%) | 1399 (51.7%) | 40.86 (10.92) | 3.65 (0.83) | 4.62 (1.04) | 0.794 (0.07) |
| The Cardiovascular Risk in Young Finns Study (YFS) | 434 | 198 (47.3%) | 186 (44.4%) | 38.88 (5.07) | 3.73 (0.75) | 4.68 (0.99) | 0.800 (0.06) |
| Finnish Twin Cohort (FTC) | 214 | 0 (0%) | 0 (0%) | 68.73 (3.31) | 2.18 (0.47) | 2.79 (0.58) | 0.786 (0.08) |
| **Total** | **23,398** |  |  |  |  |  |  |
| CHARGE Studies (European Ancestry) | Total Sample | n (%) Male | Ever Smokers, n (%) | Age, mean (SD) | FEV<sub>1</sub>, litres. mean (SD) | FVC, litres. mean (SD) | FEV<sub>1</sub>/FVC, mean (SD) |
| AGES-Reykjavik study (AGES) | 1566 | 649 (41.4%) | 900 (57.5%) | 76.1 (5.62) | 2.13 (0.70) | 2.87 (0.86) | 0.744 (0.09) |
| Atherosclerosis Risk in Communities Study (ARIC) | 10,680 | 5015 (47.0%) | 631 (59.1%) | 54.3 (5.70) | 2.94 (0.77) | 3.98 (0.98) | 0.738 (0.07) |
| Cardiovascular Health Study (CHS) | 3967 | 1737 (43.8%) | 2089 (52.7%) | 72.8 (5.55) | 2.11 (0.66) | 3.00 (0.86) | 0.702 (0.10) |
| NHLBI Family Heart Study (FAMHS) | 1651 | 718 (43.5%) | 698 (42.3) | 53.5 (12.60) | 2.91 (0.853) | 3.89 (1.05) | 0.746 (0.08) |
| Framingham Heart Study (FHS) | 7113 | 3241 (45.5%) | 3780 (53.1) | 50.7 (14.12) | 3.10 (0.925) | 4.09 (1.12) | 0.755 (0.08) |
| Health Aging and Body Composition Study (HABC) | 1457 | 786 (53.2%) | 831 (56.5%) | 73.7 (2.83) | 2.31 (0.66) | 3.11 (0.81) | 0.741 (0.08) |
| Health2006 Study | 2714 | 1217 (44·8%) | 1577 (58·1%) | 49·4 (13·04) | 3·13 (0·82) | 3·99 (0·99) | 0·784 (0·07) |
| Health2008 Study | 687 | 297 (43·2%) | 384 (55·9%) | 46·7 (8·22) | 3·27 (0·79) | 4·13 (0·97) | 0·791 (0·06) |
| Inter99 Study (without pack-years) | 1115 | 549 (49·2%) | 1115 (100%) | 47·2 (7·76) | 3·26 (0·71) | 4·12 (0·92) | 0·796 (0·07) |
| Inter99 Study (with pack-years) | 4179 | 2027 (48·5%) | 2307 (55·2%) | 45·8 (7·95) | 3·21 (0·76) | 4·10 (0·97) | 0·788 (0·08) |
| Multi-Ethnic Study of Atherosclerosis (MESA) | 1323 | 654 (49·4%) | 751 (56·8%) | 66·0 (9·8) | 2·57 (0·76) | 3·51 (0·10) | 0·733 (0·08) |
| The Rotterdam Study (RS) | 546 | 299 (54·8%) | 382 (70·0%) | 79·4 (5·00) | 2·27 (0·68) | 3·03 (0·86) | 0·750 (0·08) |
| **Total** | **36,998** |  |  |  |  |  |  |
| **CHARGE Studies (African Ancestry)** | **Total Sample** | **n (%) Male** | **Ever Smokers, n (%)** | **Age, mean (SD)** | **FEV<sub>1</sub>, litres. mean (SD)** | **FVC, litres. mean (SD)** | **FEV<sub>1</sub>/FVC, mean (SD)** |
| Atherosclerosis Risk in Communities Study (ARIC) | 3180 | 1183 (37·2%) | 1680 (59·1%) | 53·6 (5·83) | 2·48 (0·65) | 3·25 (0·82) | 0·765 (0·08) |
| Cardiovascular Health Study (CHS) | 624 | 232 (37·2%) | 340 (54·4%) | 73·2 (5·49) | 1·76 (0·58) | 2·48 (0·80) | 0·717 (0·11) |
| Health Aging and Body Composition Study (HABC) | 943 | 433 (45·9%) | 543 (57·6%) | 73·4 (2·90) | 1·96 (0·57) | 2·61 (0·71) | 0·749 (0·09) |
| Jackson Heart Study (JHS) | 2143 | 793 (36·8%) | 688 (31·9%) | 52·8 (12·6) | 2·43 (0·72) | 3·02 (0·86) | 0·807 (0·09) |
| Multi-Ethnic Study of Atherosclerosis (MESA) | 861 | 404 (46·9%) | 467 (54·2%) | 65·6 (9·6) | 2·19 (0·66) | 2·92 (0·86) | 0·756 (0·09) |
| **Total** | **7721** |  |  |  |  |  |  |
| **Replication Studies** |  |  |  |  |  |  |  |
| **Study Name** | **Total Sample** | **n (%) Male** | **Ever Smokers, n (%)** | **Age, mean(SD)** | **FEV<sub>1</sub>, litres. mean (SD)** | **FVC, litres. mean (SD)** | **FEV<sub>1</sub>/FVC, mean (SD)** |
| UK Biobank | 98,657 | 45,166 (45·8%) | 56,404 (57·2%) | 56·7 (7·92) | 2·75 (0·80) | 3·67 (0·98) | 0·75 (0·07) |
| UK Household Longitudinal Study (UKHLS) | 7443 | 3293 (44·2%) | 4509 (60·5%) | 53·10 (15·94) | 2·89 (0·90) | 3·83 (1·08) | 0·753 (0·09) |
| Netherlands Epidemiology of Obesity study (NEO) | 5456 | 2672 (48·0%) | 3674 (66·0%) | 55·9 (5·9) | 3·26 (0·80) | 4·26 (1·02) | 0·77 (0·07) |
| **Total** | **111,556** |  |  |  |  |  |  |

### Meta-analyses of single variant associations

We first evaluated single variant associations between FEV1, FVC and FEV1/FVC and the 179,215 SNPs which passed study level quality control and were polymorphic in both consortia. These analyses identified 34 SNPs in regions not previously associated with lung function, showing association with at least one trait at overall P<10^-5^, and showing association with consistent direction and P<0·05 in both consortia (full results in Supplementary Table 2, quantile-quantile and Manhattan plots shown in Supplementary Figure 1). We followed up these SNP associations in a replication analysis comprising 3 studies with 111,556 individuals. Combining the results from the discovery and replication stages in a meta-analysis identified six SNPs in total that were independent to known signals and met the pre-defined significance threshold (P<2·8x10^-7^) overall in, or near to *FGF10*, *LY86*, *SEC24C*, *RPAP1, CASC17* and *UQCC1* (Table 2, Supplementary Figure 2). A SNP near to the *CASC17* signal (rs11654749, r^2^=0·3 with rs1859962) has previously been associated with FEV1 in a genome-wide analysis of gene-smoking interactions, although this association was not replicated at the time;^20^ the present analysis provides the first evidence for independent replication of this signal. A seventh signal was also identified in *LCT* (Table 2, Supplementary Figure 2); whilst this locus has not previously been implicated in lung function, this SNP is known to vary in frequency across European populations,^21^ and we cannot rule out that this association is not an artefact of population structure. Our discovery analysis furthermore identified associations (P<10^-5^) in 25 regions previously associated with one or more of FEV1, FVC and FEV1/FVC (Supplementary Table 3).

**Table 2.**
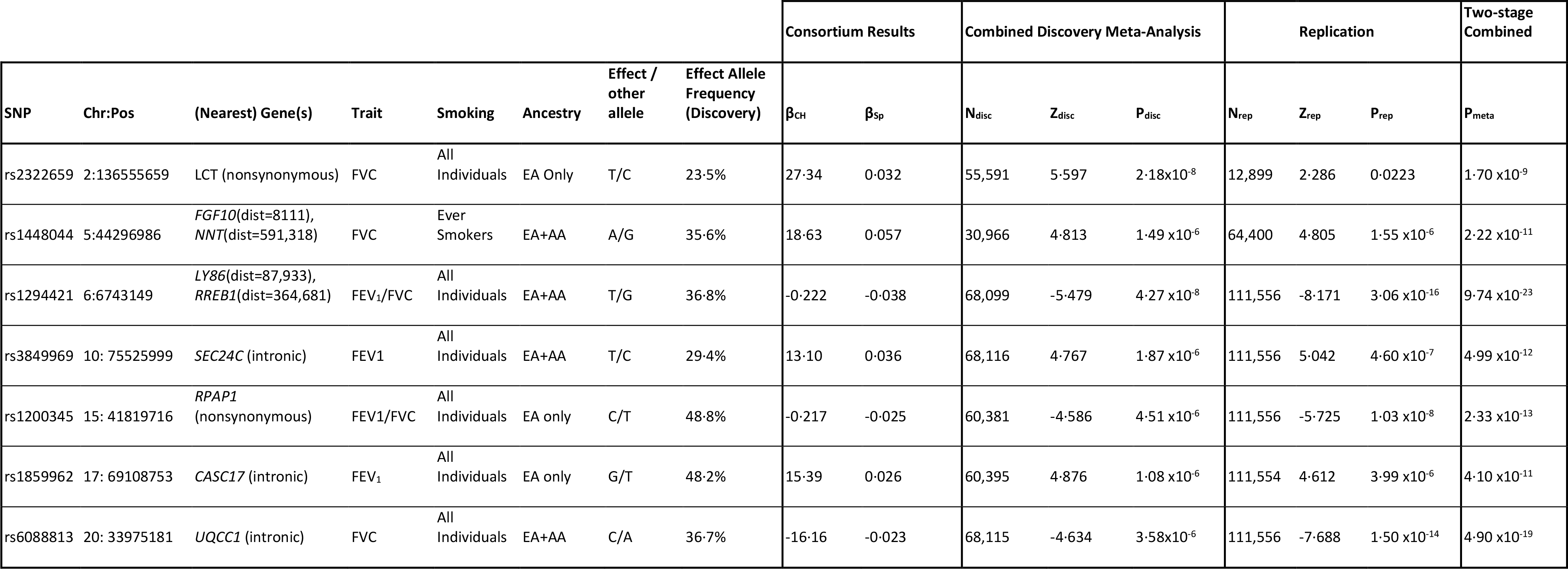
Novel loci associated with lung function traits. Results are shown for variant in novel loci associated (P<2·7×10-7) with lung function traits in a two stage meta-analysis consisting of up to 68,470 individuals from the SpiroMeta and CHARGE Consortia in the discovery analyses, with follow-up in up to 111,556 individuals from UK Biobank, UKHLS and NEO. For each SNP, the result for the trait-smoking-ancestry combination which resulted in the most statistically significant association is given. The results for these SNPs and all three traits are shown in Supplementary Table 12. Beta values from SpiroMeta (**β_Sp_**) reflect effect-size estimates on an inverse-normal transformed scale after adjustments for age, age2, sex, height and ancestry principal components, and stratified by ever smoking status (Analysis of All individuals only). Beta values from CHARGE (**βCH**) reflect effect-size estimates on an untransformed scale (litres for FEV1 and FVC; ratio for FEV1/FVC). Samples sizes (**N**), Z-statistics (**Z**) and two-sided P-values (**P**) are given for the combined discovery analysis and the replication analysis. Two-sided P-values are also given for the full two-stage combined analyses (discovery + replication).

Generally, the observed effect of the SNPs at the novel signals were similar in ever and never smokers; the exception was rs1448044 near *FGF10*, which showed a significant association with FVC only in ever smokers in our discovery analysis (ever smokers P=1·49x10^-6^; never smokers P=0·695, Supplementary Table 4 and Supplementary Figure 3). In the replication analysis, however, this association was observed in both ever and never smokers (ever smokers P=3·14x10^-5^; never smokers P=1·40x10^-4^, Supplementary Table 5). For rs1200345 (*RPAP1*) and rs1859962 (*CASC17*)), associations were most statistically significant in the analyses restricted to individuals of European Ancestry (Supplementary Table 4 and Supplementary Figure 3), as was the association with rs2322659 (*LCT*), giving further support that this association may be due to population stratification.

### Meta-Analyses of gene-based associations

We undertook Weighted Sum Tests (WST)^22^ and Sequence Kernel Association tests (SKAT)^23^ to assess the joint effects of multiple low frequency variants within genes on lung function traits. In our discovery analyses of all 68,470 individuals, we tested up to 14,380 genes that had at least two variants with MAF<5% and met the inclusion criteria (exonic or loss of function [LOF], see methods for definitions) in both consortia. The SKAT analyses identified 16 genes associated (P<0·05 in both consortia and overall P<10^-4^) with FEV1, FVC or FEV1/FVC (Supplementary Table 6), whilst the WST analyses identified 12 genes (Supplementary Table 7). There was one gene (*LY6G6D*) that was identified in both analyses. These genes were followed up in UK Biobank, with two genes, *GPR126* and *LTBP4*, showing evidence of replication in the exonic SKAT analysis (P<3·5x10^-6^); however conditional analyses in UK Biobank showed that both these associations were driven by single SNPs, that were identified in the single variant association analyses and have been previously reported in GWAS of these traits (Tables E6 & E7).

### Functional characterization of novel loci

In order to gain further insight into the six loci identified in our analyses of single variant associations (excluding *LCT*), we employed functional annotation and assessed whether identified SNPs in these regions were associated with gene expression levels. One of the identified novel SNPs was nonsynonymous, three intronic and two were intergenic. We found evidence that three of the SNPs may be involved in cis-acting regulation of the expression of several genes in multiple tissues (Supplementary Table 8).

SNP rs1200345 in *RPAP1* is a nonysynomous variant, predicted to be deleterious by both SIFT (deleterious) and Polyphen (possibly damaging) (Supplementary Table 9); *RPAP1* is ubiquitously expressed, with high levels of protein detected in the lung (Supplementary Table 10). SNP rs1200345 or proxies (r^2^>0·8) were also found to be amongst the most strongly associated SNPs with expression levels of *RPAP1* in several tissues including lung, and with a further six genes in lung tissue (Supplementary Table 8) including *TYRO3*, one of the TAM family of receptor tyrosine kinases. *TYRO3* regulates several processes including cell survival, migration and differentiation and is highly expressed in lung macrophages (Supplementary Table 10). Evidence for associations with gene expression was found at two more of the novel signals (sentinel SNPs rs3849969 and rs6088813), implicating a further 16 genes. Of note, in blood eQTL databases, a proxy of a SNP in complete linkage disequilibrium (r^2^=1) with rs3849969 (rs3812637) was an eQTL for plasminogen activator, urokinase (*PLAU*).

## Discussion

We undertook an analysis of 68,470 individuals from 23 studies with data from the exome array and three lung function traits, following up the most significant single SNP and gene-based associations in an independent sample of up to 111,556 individuals. The combined analyses of our discovery and replication single variant associations identified six SNPs meeting the pre-defined significance threshold (P<2·8x10^-7^). The replication stage results for these six SNPs also meet Bonferroni corrected significance for independent replication (P<1·47x10^-3^, corrected for 34 SNPs being tested).

One of these SNPs is in a region that has previously been implicated in lung function (near *KCJN2/SOX9*),^20^ whilst the remaining five SNPs, although all common, have not previously been identified in other GWAS of lung function. In a recent 1000 Genomes imputed analysis of lung function (which includes some of the studies contributing to the present discovery analysis), all of these SNPs showed at least suggestive association (2·97x10^-3^>P>1·28x10^-5^) with one or more lung function trait, but none reached the required threshold (P<5x10^-6^) to be taken forward for replication in that analysis.^12^

We further identified a seventh association (P<2·8x10^-7^) with rs2322659 in *LCT*. Given SNPs in this region are known to vary in frequency across European populations, we cannot dismiss the possibility that this association may be confounded by population stratification; hence we do not report this signal as a novel lung function locus. We undertook a look-up of associations in our discovery meta-analyses of 7 loci (including LCT) that were identified as showing differences in allele frequency between individuals from different regions in the UK,^24^ and subsequently across European populations.^25^ Aside from the association between the *LCT* locus and FVC, no significant associations were observed between SNPs at these loci and any lung function trait, in either the analyses restricted to EA individuals, or in the analysis of EA and AA individuals combined (Supplementary Table 11); this suggests population structure was generally accounted for adequately in our analyses.

One of the novel signals was with a nonsynonymous SNPs: rs1200345 in *RPAP1*, which is predicted to be deleterious. This SNP and proxies with r^2^>0·8 were also associated with expression in lung tissue of seven genes, including *RPAP1* and the TAM receptor *TYRO3*. TAM receptors play a role in the inhibition of Toll-like receptors (TLRs)-mediated innate immune response by initiating the transcription of cytokine signalling genes (SOCS-1 and 3) which limit cytokine overproduction and inflammation.^26,27^ It has been shown that influenza viruses H5N1 and H7N9 can cause downregulation of Tyro3, resulting in an increased inflammatory cytokine response.^27^

Three further signals were with intronic SNPs in *SEC24C*, *CASC17*, and *UQCC1*. Two of these intronic SNPs have previously been implicated in GWAS of other traits: rs1859962 in *CASC17* with prostate cancer^28^ and rs6088813 in *UQCC1* with height.^29^ The *CASC17* locus, near *KCNJ2/SOX9* has also previously been implicated in lung function, showing significant association with FEV1 in a genome-wide analysis of gene-smoking interactions, however this association was not formally replicated.^20^ Whilst the individuals utilised in the discovery stage of this analysis overlap with those included in this previous interaction analysis, the replication stage of the present study provides the first evidence of replication for this signal in independent cohorts. In the present analysis, there was no evidence that the results differed by smoking status.

SNPs rs6088813 in *UQCC1* and rs3849969 in *SEC24C* were identified as eQTLs for multiple genes. Notably, a SNP in complete linkage disequilibrium with rs3849969 (rs3812637, r^2^=1) is associated with expression of *PLAU* in blood. The plasminogen activator, urokinase (PLAU) plays a role in fibrinolysis and immunity, and with its receptor (PLAUR) is involved in degradation of the extra cellular matrix, cell migration, cell adhesion and cell proliferation.^30^ A study of preterm infants with respiratory distress syndrome, a condition characterised by intra-alveolar fibrin deposition, found *PLAU* and its inhibitor *SERPINE1* to be expressed in the alveolar epithelium, and an increased ratio of *SERPINE1* to *PLAU* was associated with severity of disease.^31^ Studies in mice have also shown that increased expression of *PLAU* may be protective against lung injury, by reducing fibrosis.^32^ *PLAU* has also been found to be upregulated in lung epithelial cells subjected to cyclic strain^33^ and in patients with COPD and lung cancer, PLAU was found to be expressed in alveolar macrophages and epithelial cells.^30^

The final two signals were with common intergenic SNPs close to *LY86* and *FGF10*. *LY86* (lymphocyte antigen 86) interacts with the Toll-like receptor signalling pathway, when bound with RP105 to form a heterodimer. ^34^ The sentinel SNP rs1294421 has previously been associated with waist-hip ratio,^35^ and an intronic SNP within *LY86* (rs7440529, LD with rs1294421: r^2^=0·005) has previously been implicated in asthma in two studies of individuals of Han Chinese ancestry. ^36,37^ FGF10 is a member of the fibroblast growth factor family of proteins, involved in a number of biological processes, including embryonic development, cell growth, morphogenesis, tissue repair, tumor growth and invasion. Specifically, the FGF10 signalling pathway plays an essential role in lung development and lung epithelial renewal.^38^ A study in mice demonstrated that a deficiency in *FGF10* resulted in a fatal disruption of branching morphogenesis during lung development.^39^

Our discovery analyses included individuals of both European and African ancestry. Two of the identified six novel signals showed inconsistent effects in the African and European ancestry individuals. For these SNPs, the associations in African Ancestry individuals were not statistically significant, and we report associations from the analysis restricted to European ancestry individuals only. For the remaining four SNPs similar effects were observed in both the European and African ancestry individuals (Supplementary Figure 3). We also examined the effects of the novel SNPs in ever smokers and never smokers separately and found these to be broadly similar, with the exception of rs1448044 in *FGF10*, which in the discovery analysis showed significant association with FVC in ever smokers, whilst showing no association in never smokers (P=0·695). In our replication stage analyses, similar effects were seen in both ever and never smokers for this SNP however, and the combined analysis of discovery and replication stages for this SNP, including both ever and never smokers, met the exome chip-wide significance level overall (P=4·22x10^-9^). We also considered whether this signal could be driven by smoking behaviour in our discovery stage as our primary analyses in SpiroMeta did not adjust for smoking quantity. We undertook a look-up of this SNP in the publicly available results of a GWAS of several smoking behaviour traits;^40^ there was only weak evidence that this SNP was associated with ever versus never smoking (P=0·039), and no evidence for association with amount smoked (cigarettes per day, P=0·10).

Through the use of the exome array, we aimed to identify associations with low frequency and rare functional variants, thereby potentially uncovering some of the missing heritability of lung function. However, whilst our discovery analyses identified single SNP associations with 23 low frequency variants (Supplementary Table 2), we did not replicate any of these findings. Eleven of these 23 SNPs we were unable to follow-up in our replication studies, due to them either being not genotyped, or monomorphic. Overall, our lack of convincing associations with rare variants is likely due to limited statistical power for identifying single variant associations, particularly if those variants exhibit only modest effects.^41^ We additionally employed SKAT and WST gene-based tests to investigate the joint effects of low frequency and rare variants within genes, on lung function traits. These analyses identified associations with a number of genes that did not appear to be driven by single SNPs. Replication of these signals proved difficult however, as again many SNPs included within the discovery stage of these analyses were not genotyped, or were monomorphic in the replication studies. This often meant a disparity in the gene unit being tested in our discovery and replication samples; hence the interpretation of these results was not straightforward. In the end, we were able to replicate only findings with common SNPs. This finding is in line with several other studies of complex traits and exome array data that have been unable to report robust associations with low frequency variants^42–44^ and it is clear that future studies, will require increasingly larger sample sizes in order to fully evaluate the effect of variants across the allele frequency spectrum. The identification of common SNPs remains important however, as such findings have the potential to highlight drug targets,^45^ and these variants collectively could have utility in risk prediction.

This study has identified six common SNPs, independent to signals previously implicated in lung function. Further interrogation of these loci could lead to greater understanding of lung function and lung disease, and could provide novel targets for therapeutic interventions.

## Methods

### Study Design, cohorts and genotyping

The SpiroMeta analysis included 23,751 individuals of European ancestry (EA) from 11 studies, and the CHARGE analysis comprised 36,998 EA individuals and a further 7,721 individuals of African ancestry (AA) from 12 studies. Follow-up analyses were conducted in an independent sample of up to 111,556 individuals from UK Biobank, the UK Household Longitudinal Study (UKHLS) and the Netherlands Epidemiology of Obesity (NEO) Study (Figure 1). All studies (excluding UK Biobank) were genotyped using either the Illumina Human Exome BeadChip v1or the Illumina Infinium HumanCoreExome-12 v1·0 BeadChip. UK Biobank samples were genotyped using the Affymetrix Axiom UK BiLEVE or UK Biobank arrays.

**Figure 1.**
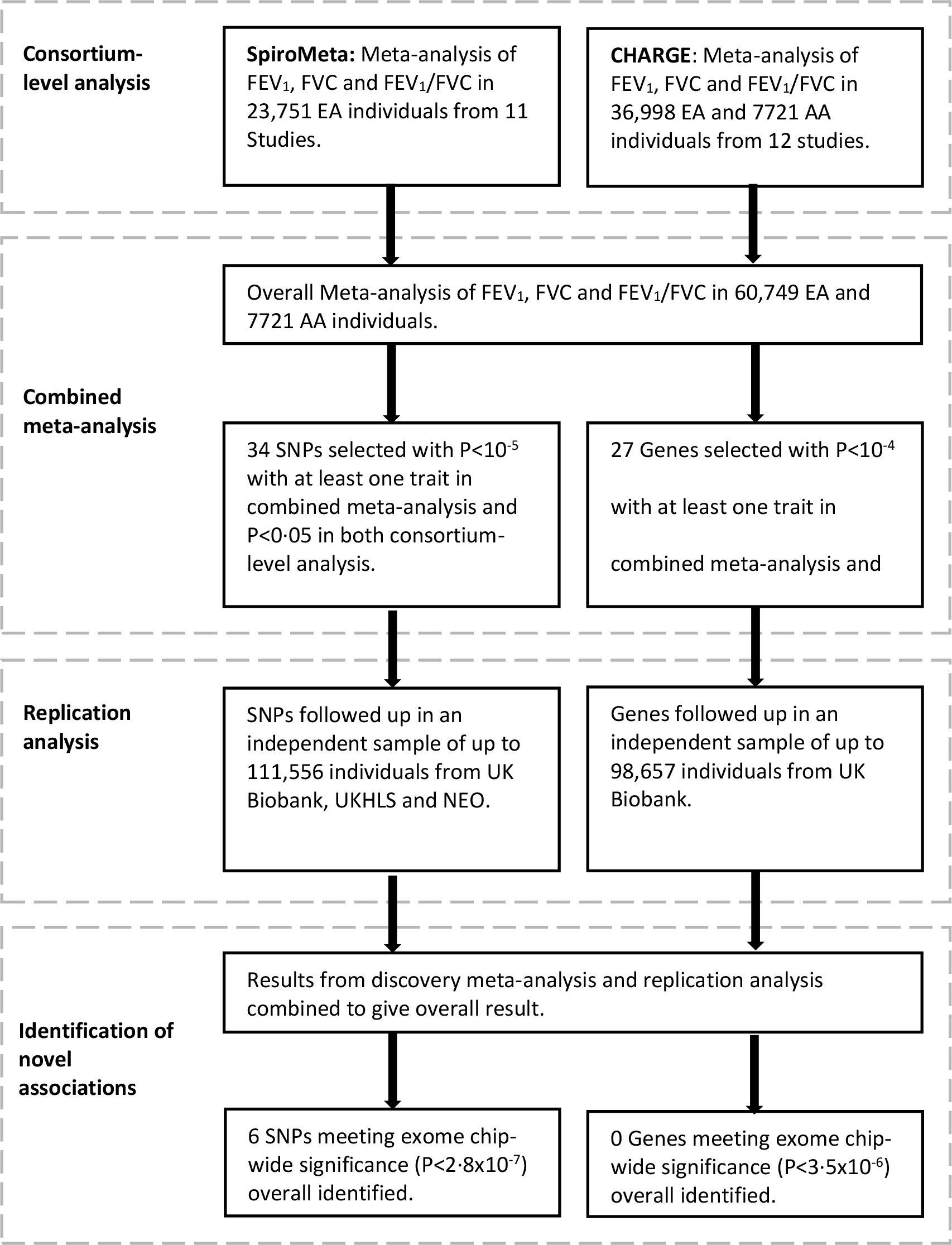
Study Design.

### Statistical analyses

#### Consortium level analyses

Within the SpiroMeta Consortium, each study contributing to the discovery analyses calculated single-variant score statistics, along with covariance matrices describing correlations between variants, using RAREMETALWORKER^46^ or rvtests.^47^ For each trait, these summary statistics were generated separately in ever and never smokers, with adjustment for sex, age, age^2^ and height, and with each trait being inverse normally transformed prior to association testing. For studies with unrelated individuals, further adjustments were made for the first 10 ancestry principal components, whilst studies with related individuals utilised linear mixed models to account for familial relationships and underlying population structure.

Within the CHARGE Consortium, each study generated equivalent summary statistics using the R package SeqMeta.^48^ For each trait, summary statistics were generated in ever and never smokers separately, and in all individuals combined. The untransformed traits were used for all analyses, adjusted for smoking status and pack-years, age, age^2^, sex, height, height^2^, centre/cohort. Models for FVC were additionally adjusted for weight. Linear regression models, with adjustment for principal components of ancestry were used for studies with unrelated individuals, and linear mixed models were used for family-based studies.

Within each consortium we used the score statistics and variance-covariance matrices generated by each study to construct both single variant and gene-based tests using either RAREMETAL^46^ (SpiroMeta) or SeqMeta^48^ (CHARGE). For single variant associations, score statistics were combined in fixed effects meta-analyses. Two gene-based tests were constructed: first, the Weighted Sum Test (WST) using Madsen Browning weightings,^22^ and secondly, the Sequence Kernel Association Test (SKAT).^23^ We performed the SKAT and WST tests using two subsets of SNPs: 1) including all SNPs with an overall consortium-wide MAF<5% that were annotated as splicing, stopgain, stoploss, or frameshift (loss of function [LOF] analysis), and 2) including all SNPs meeting the LOF analysis criteria in addition to all other nonsynonymous variants with consortium wide MAF<5% (exonic analysis). Variants were annotated to genes using dbNSFP v2·6^49^ on the basis of the GRCh37/hg19 database.

For both single variant and gene-based associations, consortium-level results were generated for ever smokers and never smokers separately, and in all individuals combined. Within the CHARGE Consortium, results were combined separately for the EA and AA studies and also in a trans-ethnic analysis of both ancestries.

#### Combined Meta-Analysis

The single variant association results from the SpiroMeta and CHARGE consortia were combined as follows: The genomic inflation statistic (λ) was calculated for SNPs with consortium-wide MAF>1%; where λ had a value greater than one, genomic control adjustment was applied to the consortium level P-values. The consortium-level results were then combined using sample size weighted z-score meta-analysis. The λ was again calculated for the meta-analysis results and genomic control applied, as appropriate. Since we were interested in identifying low frequency and rare variants, we applied no MAF or minor allele count (MAC) filter. We identified SNPs of interest as those with an overall P<10^-5^ and a consistent direction of effect and P<0·05 observed in both consortia. Where we identified a SNP within 1Mb of a previously identified lung function SNP, we deemed the SNP to represent an independent signal if it had r^2^<0·2 with the known SNP, and if it retained a P <10^-5^, when conditional analyses were carried out with the known SNP, or a genotyped proxy, using data from the SpiroMeta Consortium, or UK Biobank. Our primary meta-analysis included all individuals; we additionally carried out analyses in smoking subgroups (ever and never smokers), and in the subgroup of individuals of European ancestry only.

For genes which contained at least 2 polymorphic SNPs in both consortia, we combined the results of the consortium level gene based tests using either z-score meta-analysis (WST) or Fisher’s Method for combining P-values (SKAT). We identified genes of interest as those with P<0·05 observed in both consortia and an overall P<10^-4^. As in the analyses of single variant associations, our primary meta-analyses included all individuals, with secondary analyses undertaken in smoking and ancestry specific subgroups.

#### Replication Analyses

All SNP and gene-based associations were followed up for the trait with which they showed the most statistically significant association only. For associations identified through the smoking subgroup analyses, we followed up associations in the appropriate smoking strata; however no ancestry stratified follow-up was undertaken as replication studies included only a sufficient number of individuals of European Ancestry.

Single variant associations in UK Biobank were tested in ever smokers and never smokers separately using the score test as implemented in SNPTEST v2·5b4.^50^ Traits were adjusted for age, age^2^, height, sex, ten principal components and pack-years (ever smokers only), and inverse normally transformed. For UKHLS, analyses were undertaken analogously to the SpiroMeta discovery studies using RAREMETALWORKER, while for NEO, analyses were undertaken in the same way as was done in the CHARGE discovery studies using SeqMeta. The single variant results from all replication studies were combined using sample size weighted Z-score meta-analysis. Subsequently we combined the results from the discovery and replication stage analyses and we report SNPs with overall exome-wide significance of P<2·8x10^-7^ (Bonferroni corrected for the original 179,215 SNPs tested).

We followed up genes of interest (P<10^-4^) using data from UK Biobank only. Summary statistics for UK Biobank were generated using RAREMETALWORKER, with gene-based tests then constructed using RAREMETAL. Finally, we combined the results from the discovery analysis with the replication results in an overall combined meta-analysis using either z-score meta-analysis (WST) or Fisher’s Method (SKAT). We declared genes with overall P<3·5x10^-6^ (Bonferroni corrected for 14,380 genes tested) in our combined meta-analysis to be statistically significant. For these statistically significant genes, we carried out additional analyses using the UK Biobank data in which we conditioned on the most significantly associated individual SNP within that gene, to determine whether this was a true gene-based signal, or whether the association could be ascribed to the single SNP (if the conditional P<0·01, then association was deemed to not be driven by the single SNP).

### Characterization of findings

In order to gain further insight into the loci identified in our analyses of single variant associations, we assessed whether these regions were associated with gene expression levels in various tissues (FDR of 5%, or q-value<0·05), by querying a publically available blood eQTL database ^51^ and the GTEx project ^52^ for the sentinel SNPs, or any proxy (r^2^>0·8). We further assessed SNPs of interest (and proxies) within a lung eQTL resource based on non-tumour lung tissues of 1,111 individuals.^53–55^ Descriptions of these resources and further details of the look-ups are provided in the Supplementary Methods. Moreover, all sentinel SNPs and proxies with r^2^>0.8 were annotated using ENSEMBL’s Variant Effect Predictor (VEP);^56^ potentially deleterious coding variants were identified as those annotated as ‘deleterious’ by SIFT^57^ or ‘probably damaging’ or ‘possibly damaging’ by PolyPhen-2.^58^ For all genes implicated through the expression data or functional annotation, we searched for evidence of protein expression in the respiratory system by querying the Human Protein Atlas.^59^

## Acknowledgements

This research has been conducted using the UK Biobank Resource. This article presents independent research funded partially by the National Institute for Health Research (NIHR). The views expressed are those of the author(s) and not necessarily those of the NHS, the NIHR or the Department of Health. These data are from Understanding Society: The UK Household Longitudinal Study (UKHLS), which is led by the Institute for Social and Economic Research at the University of Essex and funded by the Economic and Social Research Council. The data were collected by NatCen and the genome wide scan data were analysed by the Wellcome Trust Sanger Institute. Information on how to access the data can be found on the Understanding Society website https://www.understandingsociety.ac.uk/. The authors would like to thank the staff at the Quebec Respiratory Health Network Tissue Bank for their valuable assistance with the lung eQTL dataset at Laval University. The content is solely the responsibility of the authors and does not necessarily represent the official views of the National Institutes of Health. A full list of principal CHS investigators and institutions can be found at CHS-NHLBI.org. The authors thank the staff and participants of the ARIC study for their important contributions. The authors of the NEO study thank all individuals who participated in the Netherlands Epidemiology in Obesity study, all participating general practitioners for inviting eligible participants and all research nurses for collection of the data. We thank the NEO study group, Pat van Beelen, Petra Noordijk and Ingeborg de Jonge for the coordination, lab and data management of the NEO study. SAPALDIA could not have been done without the help of the study participants, technical and administrative support and the medical teams and field workers at the local study sites. Local fieldworkers : Aarau: M Broglie, M Bünter, D Gashi, Basel: R Armbruster, T Damm, U Egermann, M Gut, L Maier, A Vögelin, L Walter, Davos: D Jud, N Lutz, Geneva: M Ares, M Bennour, B Galobardes, E Namer, Lugano: B Baumberger, S Boccia Soldati, E Gehrig-Van Essen, S Ronchetto, Montana: C Bonvin, C Burrus, Payerne: S Blanc, AV Ebinger, ML Fragnière, J Jordan, Wald: R Gimmi, N Kourkoulos, U Schafroth. Administrative staff: N Bauer, D Baehler, C Gabriel, R Gutknecht. SAPALDIA Team: *Study directorate:* NM Probst Hensch, T Rochat, N Künzli, C Schindler, JM Gaspoz. *Scientific team:* JC Barthélémy, W Berger, R Bettschart, A Bircher, G Bolognini, O Brändli, C Brombach, M Brutsche, L Burdet, M Frey, U Frey, MW Gerbase, D Gold, E de Groot, W Karrer, R Keller, B Knöpfli, B Martin, D Miedinger, U Neu, L Nicod, M Pons, F Roche, T Rothe, E Russi, P Schmid-Grendelmeyer, A Schmidt-Trucksäss, A Turk, J Schwartz, D. Stolz, P Straehl, JM Tschopp, A von Eckardstein, E Zemp Stutz. *Scientific team at coordinating centers:* M Adam, E Boes, PO Bridevaux, D Carballo, E Corradi, I Curjuric, J Dratva, A Di Pasquale, L Grize, D Keidel, S Kriemler, A Kumar, M Imboden, N Maire, A Mehta, F Meier, H Phuleria, E Schaffner, GA Thun, A Ineichen, M Ragettli, M Ritter, T Schikowski, G Stern, M Tarantino, M Tsai, M Wanner. MDT has been supported by MRC fellowships G0501942 and G0902313. MDT and LVW are supported by the MRC (MR/N011317/1). IPH is supported by the MRC (G1000861). ALW and SJL are supported by the Intramural Research Program of the NIH, National Institute of Environmental Health Sciences (ZIA ES 043012). We acknowledge use of phenotype and genotype data from the British 1958 Birth Cohort DNA collection, funded by the Medical Research Council grant G0000934 and the Wellcome Trust grant 068545/Z/02. APM is a Wellcome Trust Senior Fellow in Basic Biomedical Science (grant number WT098017) and also supported by Wellcome Trust grant WT064890. EI is supported by the Swedish Research Council (2012-1397), Knut och Alice Wallenberg Foundation (2013.0126) and the Swedish Heart-Lung Foundation (20140422). JK is supported by Academy of Finland Center of Excellence in Complex Disease Genetics grants 213506, 129680 and Academy of Finland grants 265240, 263278. The Finnish Twin Cohort is supported by the Welcome Trust Sanger Institute, UK. The Lothian Birth Cohort is supported by Age UK (The Disconnected Mind Project), the UK Medical Research Council (MR/K026992/1) and The Royal Society of Edinburgh. ÅJ is supported by the Swedish Society for Medical Research (SSMF), The Kjell och Märta Beijers Foundation, The Marcus Borgström Foundation, The Åke Wiberg foundation and The Vleugels Foundation. UG is supported by Swedish Medical Research Council grants K2007-66X-20270-01-3 and 2011-2354 and European Commission FP6 (LSHG-CT-2006-01947). SHIP is part of the Community Medicine Research net of the University of Greifswald, Germany, which is funded by the Federal Ministry of Education and Research, the Ministry of Cultural Affairs as well as the Social Ministry of the Federal State of Mecklenburg-West Pomerania, and the network ‘Greifswald Approach to Individualized Medicine (GANI_MED)’ funded by the Federal Ministry of Education and Research, and the German Asthma and COPD Network (COSYCONET) (grant no.01ZZ9603, 01ZZ0103, 01ZZ0403, 03IS2061A, BMBF 01GI0883). ExomeChip data have been supported by the Federal Ministry of Education and Research (grant no. 03Z1CN22) and the Federal State of Mecklenburg-West Pomerania. The University of Greifswald is a member of the Caché Campus program of the InterSystems GmbH. UKHLS is supported by grants WT098051 (Wellcome Trust) and ES/H029745/1 (Economic and Social Research Council). Y.B. holds a Canada Research Chair in Genomics of Heart and Lung Diseases. Lies Lahousse is a Postdoctoral Fellow of the Research Foundation - Flanders (FWO grant G035014N). The Rotterdam Study is funded by Erasmus Medical Center and Erasmus University, Rotterdam, the Netherlands Organization for Scientific Research (NOW), the Netherlands Organization for the Health Research and Development (ZonMw), the Research Institute for Diseases in the Elderly (RIDE), the Ministry of Education, Culture and Science, the Ministry for Health, Welfare and Sports, the European Commission (DG XII), and the Municipality of Rotterdam. Genotyping in the Rotterdam study was supported by Netherlands Organization for Scientific Research (NOW grants 175.010.2005.011; 911-03-305 012), the Research Institute for Diseases in the Elderly (RIDE2 grants 014-93-015) and Netherlands Genomics Initiative (NGI)/Netherlands Consortium for Healthy Aging (NCHA grant050-060-810). MESA/MESA SHARe is supported by HHS (HHSN268201500003I), NIH/NHLBI (contracts N01-HC-95159, N01-HC-95160, N01-HC-95161, N01-HC-95162, N01-HC-95163, N01-HC-95164, N01-HC-95165, N01-HC-95166, N01-HC-95167, N01-HC-95168, N01-HC-95169) and HIH/NCATS (contracts UL1-TR-000040, UL1-TR-001079, UL1-TR-001881, DK063491). MESA SHARe is funded by NIH/NHLBI contract N02-HL-64278, MESA Air is funded by US EPA (RD831697) and MESA Spirometry funded by NIH/NHLBI (R01-HL077612). SSR and BMP are supported by NIH/NHLBI grant rare variants and NHLBI traits in deeply phenotyped cohorts (R01-HL120393). Cardiovascular Health Study: This CHS research was supported by NHLBI contracts HHSN268201200036C, HHSN268200800007C, N01HC55222, N01HC85079, N01HC85080, N01HC85081, N01HC85082, N01HC85083, N01HC85086; and NHLBI grants U01HL080295, R01HL068986, R01HL087652, R01HL105756, R01HL103612, R01HL120393, and R01HL130114 with additional contribution from the National Institute of Neurological Disorders and Stroke (NINDS). Additional support was provided through R01AG023629 from the National Institute on Aging (NIA). The provision of genotyping data was supported in part by the National Center for Advancing Translational Sciences, CTSI grant UL1TR000124, and the National Institute of Diabetes and Digestive and Kidney Disease Diabetes Research Center (DRC) grant DK063491 to the Southern California Diabetes Endocrinology Research Center. The Atherosclerosis Risk in Communities (ARIC) study is carried out as a collaborative study supported by the National Heart, Lung, and Blood Institute (NHLBI) contracts (HHSN268201100005C, HHSN268201100006C, HHSN268201100007C, HHSN268201100008C, HHSN268201100009C, HHSN268201100010C, HHSN268201100011C, and HHSN268201100012C). Funding support for “Building on GWAS for NHLBI-diseases: the U.S. CHARGE consortium” was provided by the NIH through the American Recovery and Reinvestment Act of 2009 (ARRA) (5RC2HL102419). DOMK received funding from the Dutch Science Organisation (ZonMW-VENI Grant 916.14.023). The genotyping in the NEO study was supported by the Centre National de Génotypage (Paris, France), headed by Jean-François Deleuze. The NEO study is supported by the participating Departments, the Division and the Board of Directors of the Leiden University Medical Center, and by the Leiden University, Research Profile Area Vascular and Regenerative Medicine. SAPALDIA was supported by the Swiss National Science Foundation (grants no 33CS30-148470/1, 33CSCO- 134276/1, 33CSCO-108796, 324730_135673, 3247BO-104283, 3247BO-104288, 3247BO-104284, 3247-065896, 3100-059302, 3200-052720, 3200-042532, 4026-028099, PMPDP3_129021/1, PMPDP3_141671/1), the Federal Office for the Environment, the Federal Office of Public Health, the Federal Office of Roads and Transport, the canton’s government of Aargau, Basel-Stadt, Basel-Land, Geneva, Luzern, Ticino, Valais, and Zürich, the Swiss Lung League, the canton’s Lung League of Basel Stadt/Basel Landschaft, Geneva, Ticino, Valais, Graubünden and Zurich, Stiftung ehemals Bündner Heilstätten, SUVA, Freiwillige Akademische Gesellschaft, UBS Wealth Foundation, Talecris Biotherapeutics GmbH, Abbott Diagnostics, European Commission 018996 (GABRIEL), Wellcome Trust WT 084703MA. The Novo Nordisk Foundation Center for Basic Metabolic Research is an independent Research Center at the University of Copenhagen partially funded by an unrestricted donation from the Novo Nordisk Foundation (www.metabol.ku.dk). Generation Scotland received core support from the Chief Scientist Office of the Scottish Government Health Directorates [CZD/16/6] and the Scottish Funding Council [HR03006]. Genotyping of the GS:SFHS samples was carried out by the Genetics Core Laboratory at the Edinburgh Clinical Research Facility, University of Edinburgh, Scotland and was funded by the Medical Research Council UK. The Croatia KORCULA study was supported by the Ministry of Science, Education and Sport in the Republic of Croatia (108-1080315-0302). JD, JCL, WG and GTOC are supported by NIH/NHLBI Contract HHSN268201500001I. Genotyping, quality control and calling of the Illumina HumanExome BeadChip in the Framingham Heart Study was supported by funding from the National Heart, Lung and Blood Institute Division of Intramural Research (Daniel Levy and Christopher J. O’Donnell, Principle Investigators). The AGES study is supported by the NIH (N01-AG012100), the Iceland Parliament (Alþingi) and the Icelandic Heart Association. HABC was supported by NIA contracts N01AG62101, N01AG62103, and N01AG62106; NIA grant R01-AG028050, and NINR grant R01- NR012459 and was supported in part by the Intramural Research Program of the NIH, National Institute on Aging. The HABC genome-wide association study was funded by NIA grant 1R01AG032098- 01A1 and genotyping services were provided by the Center for Inherited Disease Research (CIDR). CIDR is fully funded through a federal contract from the National Institutes of Health to The Johns Hopkins University, contract number HHSN268200782096C.

This research used the ALICE and SPECTRE High Performance Computing Facilities at the University of Leicester.

## Author Contributions

Ordered Alphabetically:

ABW, AGE, AL, BMP, BS, CH, CP, DOMK, DPS, EZ, GGB, HS, IPH, JBJ, JK, KMB, LL, MAI, MAP, MDT, MK, NG, NMPH, OP, OTR, RdM, RGB, SBK, SG, SJL, SSR, TA, TBH, TH, TL, TR, TS, UG contributed to study concept and designs. AC, AJ, A.Manichaikul, BHS, BMP, BS, CP, DJP, DPS, EI, GGB, GTOC, IJD, JBJ, JGW, JK, JMS, KS, LAL, LL, LL, MAP, MI, MK, NG, NMPH, OP, OTR, PAC, RdM, RGB, RR, SBK, SE, SEH, SG, SK, SK, TA, TBH, TDP, TL, TNB, TR, UG, WT, WT contributed to phenotype data acquisition and quality control. AGE, AJ, AK, AK, ALT, ALT, A.Manichaikul, APM, AT, BMP, BP, CH, DOMK, EI, GD, HV, IJD, JAB, JCM, JGW, JL, KDT, KEN, KL, L-PL, LAL, LL, MAP, MI, MLG, NMPH, OP, RGB, RLG, RR, SBK, SE, SEH, SRH, SSR, SW, TBH, TDP, TH, TL, YL contributed to genotype data acquisition and quality control. DDS, KH, WT, YB contribute to eQTL data acquisition and quality control. ABW, ACM, AK, AK, ALT, A.Mahajan, A.Manichaikul, APM, AT, BP, BQ, CH, CMS, EA, HV, IPH, JAB, JCL, JD, JEH, JL, JM, JMJ, KL, L-PL, LL, LVW, MDT, MI, MO, NF, NMPH, OP, PAC, RLG, SE, SEH, SJL, SW, TDP, TH, TMB, VEJ, WG, WT, YL contributed to data analysis. All authors contributed to writing and/or critical review of the manuscript.

The ‘Understanding Society Scientific Group’ include the following: Understanding Society Scientific Group: Michaela Benzeval, Jonathan Burton, Nicholas Buck, Annette Jäckle, Meena Kumari, Heather Laurie, Peter Lynn, Stephen Pudney, Birgitta Rabe, Shamit Saggar, Noah Uhrig, Dieter Wolke.

## References

1. Rabe KF, Hurd S, Anzueto A, et al. Global Strategy for the Diagnosis, Management, and Prevention of Chronic Obstructive Pulmonary Disease. Am J Respir Crit Care Med 2007; 176(6): 532–55.

2. Palmer LJ, Knuiman MW, Divitini ML, et al. Familial aggregation and heritability of adult lung function: results from the Busselton Health Study. Eur Respir J 2001; 17(4): 696–702.

3. Wilk JB, DeStefano AL, Joost O, et al. Linkage and association with pulmonary function measures on chromosome 6q27 in the Framingham Heart Study. Human Molecular Genetics 2003; 12(21): 2745–51.

4. Klimentidis YC, Vazquez AI, de lC, Allison DB, Dransfield MT, Thannickal VJ. Heritability of pulmonary function estimated from pedigree and whole-genome markers. Front Genet 2013; 4: 174.

5. Wilk JB, Djousse L, Arnett DK, et al. Evidence for major genes influencing pulmonary function in the NHLBI Family Heart Study. Genet Epidemiol 2000; 19(1): 81–94.

6. Wilk JB, Chen T, Gottlieb DJ, et al. A Genome-Wide Association Study of Pulmonary Function Measures in the Framingham Heart Study. PLoS Genet 2009; 5(3): e1000429.

7. Repapi E, Sayers I, Wain LV, et al. Genome-wide association study identifies five loci associated with lung function. Nat Genet 2010; 42(1): 36–44.

8. Soler Artigas M, Loth DW, Wain LV, et al. Genome-wide association and large-scale follow up identifies 16 new loci influencing lung function. Nat Genet 2011; 43(11): 1082–90.

9. Hancock DB, Eijgelsheim M, Wilk JB, et al. Meta-analyses of genome-wide association studies identify multiple loci associated with pulmonary function. Nat Genet 2010; 42(1): 45–52.

10. Loth DW, Soler Artigas M, Gharib SA, et al. Genome-wide association analysis identifies six new loci associated with forced vital capacity. Nat Genet 2014; 46(7): 669–77.

11. Wain LV, Shrine N, Miller S, et al. Novel insights into the genetics of smoking behaviour, lung function, and chronic obstructive pulmonary disease (UK BiLEVE): a genetic association study in UK Biobank. Lancet Respir Med 2015; 3(10): 769–81.

12. Soler Artigas M, Wain LV, Miller S, et al. Sixteen new lung function signals identified through 1000 Genomes Project reference panel imputation. Nat Commun 2015; 6: 8658.

13. Wain LV, Shrine N, Soler Artigas M, et al. Genome-wide association analyses for lung function and chronic obstructive pulmonary disease identify new loci and potential druggable targets. 2017;.

14. Pillai SG, Ge D, Zhu G, et al. A Genome-Wide Association Study in Chronic Obstructive Pulmonary Disease (COPD): Identification of Two Major Susceptibility Loci. PLoS Genet 2009; 5(3): e1000421.

15. Cho MH, Boutaoui N, Klanderman BJ, et al. Variants in FAM13A are associated with chronic obstructive pulmonary disease. Nat Genet 2010; 42(3): 200–2.

16. Cho MH, McDonald MN, Zhou X, et al. Risk loci for chronic obstructive pulmonary disease: a genome-wide association study and meta-analysis. The Lancet Respiratory Medicine 2014; 2(3): 214–25.

17. Hobbs BD, de Jong K, Lamontagne M, et al. Genetic loci associated with chronic obstructive pulmonary disease overlap with loci for lung function and pulmonary fibrosis. Nat Genet 2017; 49(3): 426–32.

18. Hobbs BD, Parker MM, Chen H, et al. Exome Array Analysis Identifies A Common Variant in IL27 Associated with Chronic Obstructive Pulmonary Disease. Am J Respir Crit Care Med 2016; : doi:10.1164/rccm.201510-2053OC.

19. Abecasis GR. Exome Chip Design Wiki. 2013; Available at: http://genome.sph.umich.edu/wiki/Exome_Chip_Design. Accessed August 30, 2013.

20. Hancock DB, Soler Artigas M, Gharib SA, et al. Genome-wide joint meta-analysis of SNP and SNP-by-smoking interaction identifies novel loci for pulmonary function. PLoS Genet 2012; 8(12): e1003098.

21. Campbell CD, Ogburn EL, Lunetta KL, et al. Demonstrating stratification in a European American population. Nat Genet 2005; 37(8): 868–72.

22. Madsen BE, Browning SR. A Groupwise Association Test for Rare Mutations Using a Weighted Sum Statistic. PLoS Genet 2009; 5(2): e1000384.

23. Wu M, Lee S, Cai T, Li Y, Boehnke M, Lin X. Rare-Variant Association Testing for Sequencing Data with the Sequence Kernel Association Test. Am J Hum Genet 2011; 89(1): 82–93.

24. Burton PR, Clayton DG, Cardon LR, et al. Genome-wide association study of 14,000 cases of seven common diseases and 3,000 shared controls. Nature 2007; 447(7145): 661–78.

25. Heath SC, Gut IG, Brennan P, et al. Investigation of the fine structure of European populations with applications to disease association studies. European Journal of Human Genetics 2008; 16(12): 1413–29.

26. Grabiec AM, Hussell T. The role of airway macrophages in apoptotic cell clearance following acute and chronic lung inflammation. Seminars in immunopathology; Springer; 2016.

27. Ye S, Lowther S, Stambas J. Inhibition of reactive oxygen species production ameliorates inflammation induced by influenza A viruses via upregulation of SOCS1 and SOCS3. J Virol 2015; 89(5): 2672–83.

28. Gudmundsson J, Sulem P, Steinthorsdottir V, et al. Two variants on chromosome 17 confer prostate cancer risk, and the one in TCF2 protects against type 2 diabetes. Nat Genet 2007; 39(8): 977–83.

29. Soranzo N, Rivadeneira F, Chinappen-Horsley U, et al. Meta-analysis of genome-wide scans for human adult stature identifies novel Loci and associations with measures of skeletal frame size. PLoS Genet 2009; 5(4): e1000445.

30. Wang I, Stepaniants S, Boie Y, et al. Gene expression profiling in patients with chronic obstructive pulmonary disease and lung cancer. American journal of respiratory and critical care medicine 2008; 177(4): 402–11.

31. Cederqvist K, Sirén V, Petäjä J, Vaheri A, Haglund C, Andersson S. High concentrations of plasminogen activator inhibitor-1 in lungs of preterm infants with respiratory distress syndrome. Pediatrics 2006; 117(4): 1226–34.

32. Sisson TH, Hanson KE, Subbotina N, Patwardhan A, Hattori N, Simon RH. Inducible lung-specific urokinase expression reduces fibrosis and mortality after lung injury in mice. Am J Physiol Lung Cell Mol Physiol 2002; 283(5): L1023–32.

33. Weber B, Bader N, Lehnich H, Simm A, Silber RE, Bartling B. Microarray-based gene expression profiling suggests adaptation of lung epithelial cells subjected to chronic cyclic strain. Cell Physiol Biochem 2014; 33(5): 1452–66.

34. Kimoto M, Nagasawa K, Miyake K. Role of TLR4/MD-2 and RP105/MD-1 in innate recognition of lipopolysaccharide. Scand J Infect Dis 2003; 35(9): 568–72.

35. Heid IM, Jackson AU, Randall JC, et al. Meta-analysis identifies 13 new loci associated with waist-hip ratio and reveals sexual dimorphism in the genetic basis of fat distribution. Nat Genet 2010; 42(11): 949–60.

36. Tan JY, Luo YL, Huang X, Shao JL, Lin L, Yang XX. Association of single nucleotide polymorphisms of MD-1 gene with asthma in adults of Han Nationality in Southern China. Zhonghua Jie He He Hu Xi Za Zhi 2011; 34(2): 104–8.

37. Lee S, Wang J, Hsieh Y, Wu Y, Ting H, Wu L. Association of single nucleotide polymorphisms of MD-1 gene with pediatric and adult asthma in the Taiwanese population. J Microbiol Immunol Infect 2008; 41(6): 445–9.

38. Klar J, Blomstrand P, Brunmark C, et al. Fibroblast growth factor 10 haploinsufficiency causes chronic obstructive pulmonary disease. Journal of Medical Genetics 2011; 48(10): 705–9.

39. Sekine K, Ohuchi H, Fujiwara M, et al. Fgf10 is essential for limb and lung formation. Nat Genet 1999; 21(1): 138–41.

40. The Tobacco and Genetics Consortium. Genome-wide meta-analyses identify multiple loci associated with smoking behavior. Nat Genet 2010; 42(5): 441–7.

41. Manolio TA, Collins FS, Cox NJ, et al. Finding the missing heritability of complex diseases. Nature 2009; 461(7265): 747–53.

42. Zuo X, Sun L, Yin X, et al. Whole-exome SNP array identifies 15 new susceptibility loci for psoriasis. Nat Commun 2015; 6: 6793.

43. Holmen OL, Zhang H, Zhou W, et al. No large-effect low-frequency coding variation found for myocardial infarction. Human Molecular Genetics 2014; 23(17): 4721–8.

44. Tajuddin SM, Schick UM, Eicher JD, et al. Large-scale exome-wide association analysis identifies loci for white blood cell traits and pleiotropy with immune-mediated diseases. The American Journal of Human Genetics 2016; 99(1): 22–39.

45. Nelson MR, Tipney H, Painter JL, et al. The support of human genetic evidence for approved drug indications. Nat Genet 2015; 47(8): 856–60.

46. Liu DJ, Peloso GM, Zhan X, et al. Meta-Analysis of Gene Level Association Tests. Nat Genet 2014; 46(2): 200–4.

47. Zhan X, Hu Y, Li B, Abecasis G, Liu D. RVTESTS: an efficient and comprehensive tool for rare variant association analysis using sequence data. Bioinformatics 2016; 32(9): 1423–6.

48. Lumley T, Brody J, Dupus J, Cupples A. Meta-analysis of a rare-variant association test. 2012; Available at: http://stattech.wordpress.fos.auckland.ac.nz/files/2012/11/skat-meta-paper.pdf.

49. Liu X, Jian X, Boerwinkle E. dbNSFP v2. 0: a database of human non-synonymous SNVs and their functional predictions and annotations. Hum Mutat 2013; 34(9): E2393–402.

50. Marchini J, Howie B, Myers S, McVean G, Donnelly P. A new multipoint method for genome-wide association studies by imputation of genotypes. Nat Genet 2007; 39(7): 906–13.

51. Westra H, Peters MJ, Esko T, et al. Systematic identification of trans eQTLs as putative drivers of known disease associations. Nat Genet 2013; 45(10): 1238–43.

52. Lonsdale J, Thomas J, Salvatore M, et al. The Genotype-Tissue Expression (GTEx) project. Nat Genet 2013; 45(6): 580–5.

53. Hao K, Bossé Y, Nickle DC, et al. Lung eQTLs to help reveal the molecular underpinnings of asthma. PLoS Genet 2012; 8(11): e1003029.

54. Lamontagne M, Couture C, Postma DS, et al. Refining susceptibility loci of chronic obstructive pulmonary disease with lung eqtls. PLoS One 2013; 8(7): e70220.

55. Obeidat M, Miller S, Probert K, et al. GSTCD and INTS12 Regulation and Expression in the Human Lung. PLoS ONE 2013; 8(9): e74630.

56. McLaren W, Gil L, Hunt SE, et al. The Ensembl Variant Effect Predictor. Genome Biol 2016; 17(1): 1.

57. Kumar P, Henikoff S, Ng PC. Predicting the effects of coding non-synonymous variants on protein function using the SIFT algorithm. Nature protocols 2009; 4(7): 1073–81.

58. Adzhubei IA, Schmidt S, Peshkin L, et al. A method and server for predicting damaging missense mutations. Nature methods 2010; 7(4): 248–9.

59. Uhlen M, Oksvold P, Fagerberg L, et al. Towards a knowledge-based Human Protein Atlas. Nat Biotech 2010; 28(12): 1248–50.

